# Impact of zoonotic disease outbreaks on international wildlife trade dynamics in Ghana

**DOI:** 10.1101/2021.09.18.460894

**Authors:** Godfred Bempah, Moses A. Nartey, Kwaku B. Dakwa, Kevin Messenger

**Author notes:** Nanjing Forestry University, 159 Longpan Road, Nanjing, Jiangsu, China.

## Abstract

Wildlife is under intense pressure from trade, which most likely contributes to zoonotic diseases. The study explores the impact of zoonotic disease outbreaks on the wildlife trade of Ghana. This study provides an in-depth analysis of the trends of taxa trade and factors that influence trade from 1975–2018 by combining zoonotic disease data with data from the CITES database. Trade flow showed that reptiles were the most traded group, followed by birds, mammals, and amphibians. Species of the families Pythonidae, Dendrobatidae, Cercopithecidae, and Psittacidae were the most traded. The decade mean number of trade for 1997–2007 was the highest (n = 62) followed by 2008–2018 (n = 54.4). Most exporter countries that traded with Ghana are from Africa and importers from the United States of America, Europe and Asia. Continuous trade in reptiles and birds, especially the endangered pythons and psittacus species, could lead to their extinction in the wild. The outbreak of zoonotic diseases influenced the dynamics of the wildlife trade in Ghana as traders shifted their activities among taxa over a period of time. Because those taxa were observed to harbour zoonotic diseases and constitute high health risks when traded. Mammals’ trade flow decreased with disease outbreaks over time, while reptiles increased. Early detection of zoonotic diseases and the adoption of an expanded education module on avoiding species capable of harbouring pathogens will most likely help reduce trade in wildlife.

## Introduction

The less emphasis and publicised problems of the substantial impact of wildlife trade over the years could lead to over-exploitation and extinction of species (Shepherd and Nijman 2008). This is evident in the loss of global biodiversity through poaching, death from unwarranted concealment when transporting the animals, and the introduction of non-native species. Actors in the trade value chain experience direct or indirect forms of contact with animals resulting in over a billion multiple contacts (Karesh et al. 2005).

Several species of live and dead wild animals and associated elements are transported all over the world (Rosen and Smith 2010), with an estimated number of live species, including birds (4 million), primates (40,000), and reptiles (640,000) traded annually (World Wildlife Fund 2001). According to UNEP-WCMC (2013) CITES has compiled several millions of trade records in wildlife, with 34,000 scientific names of taxa. Legal protection of species by individual countries or international collaboration is useful in reviving the population of over-exploited species (Abotsi et al. 2016). Ghana has a rich stock of biodiversity and officially ratified the CITES agreement. Ghana falls within the Guinean Forests of West Africa Hotspot with many endemic species. This area is considered a priority for conservation at the global level against ecological destabilisers such as wildlife trade.

Aside from the ecological impact of the wildlife trade, its contribution to emerging infectious diseases is highly important (Bezerra-Santos et al. 2021). Global health for the past decade has substantially been negatively affected by infectious diseases, accounting for millions of human deaths (WHO 2018). For the past forty-one years, the world has witnessed an outbreak of over 35 infectious diseases (Karesh et al. 2005). It is estimated that about 75% of recent infectious diseases are zoonotic, with a greater proportion traced to wild animals (Jones et al. 2008). Recently, the outbreak of the Covid-19 pandemic calls for the urgent need to reconsider human-wildlife contacts (Borsky et al. 2020). Several studies have documented wildlife trade as a major source of infectious diseases (Bezerra-Santos et al. 2021; Borsky et al. 2020; Jones et al. 2008; Green et al. 2020; Karesh et al. 2005). Although characteristically, different wild animals serve as hosts for different infectious diseases (Karesh et al. 2005), it is yet not known if traders shift their trading activities among taxa groups due to a major diseases outbreak. To the best of our knowledge, there is no existence of such a trade assessment for Ghana.

Most studies concerning wildlife trade in Ghana have arisen from research about wildlife trade locally. Such studies (Gbogbo and Daniels 2019; Odonkor et al. 2007) focused on towns within Ghana. It has therefore become important to bring to light the interconnectivity of the international wildlife trade in Ghana. We seek to partially fill the knowledge gap by analysing the overall trends of species targeted over 43 years period and countries (routes) significantly involved in the trade. Second, establish if the outbreak of zoonotic disease affects taxa trade. Identifying and creating an understanding of the linkages between taxon specificity and trends will provide a scientific basis for enacting improved policy directions and approaches in tackling the associated challenges of wildlife trade. We hypothesized that; 1) there will be high trade in mammal species than other taxa, 2) Most countries in Asia will be highly involved in trade than other part of the world, and 3) Outbreak of zoonotic diseases will have a significant negative relationship with bird trade than other taxa.

## Method

### Data collection

Trade data was collected from the CITES online database managed by the World Conservation Monitoring Centre (WCMC). Records of trade activities between Ghana and other countries from 1975–2018 captured in the CITES database were extracted. Data from CITES (http://www.unep-wcmc.org/citestrade) was relied on because of its availability and also considered one of the most prominent and widely adopted mechanism internationally. The quantitative data examined for this analysis are the reported number of trade incidents.

Parameters were selected in the trade database. These were: (a) Ghana – importer country vis-a-vis all countries as exporter country, (b) Ghana – exporter country vis-a-vis all countries as importer country, (c) year range from 1975–2018. Parameters (a and c) and (b and c) were searched concurrently for each of the four taxa (amphibians, birds, mammals, and reptiles). These taxa were targets for selection because a notable feature of Ghana’s wildlife protection and utilisation policies focuses on them. For each taxon searched, we recorded the number of incidents a species was traded, the appendices, family, and the year of trade. This method was favoured because it ensures the regularity of measuring the outcomes for the various parameters (Wyatt 2016). In the case of all established trades - legal or illegal-the study acknowledges that there could be more incidents unnoticed and thus not recorded. Therefore, what has been analysed in our study are the reported incidents. Transportation of organisms harbouring pathogens is possible irrespective of the legal or illegal trade means (Can 2019); therefore, exclusive reliance on only illegal wildlife trade results in deficiency (Nijman 2021). Our study thus focused on both legal and illegal trade.

A literature search was done to identify the zoonotic pathogens known to occur using wild animals in the World Health Organization website (www.who.int), Google search engine (www.google.com), and published research. All species identified to host zoonotic pathogens were classified into four major taxa (Amphibians, birds, mammals and reptiles) and documented periods (years) of major outbreaks. The zoonotic disease was then cross-referenced with the traded (exported/imported) taxa to indicate the diseases found in each taxon for each major year of the outbreak. For a disease to be used in the analysis, it must: 1) have a record of human infection from a wild animal; 2) must have a high human health risk, e.g., death or severe illness; 3) have the year of the initial outbreak or other years of devastating effect recorded; and 4) be considered to have global or regional risk significance. The final list contained a total of 25 zoonoses of high risk.

### Data analysis

Mann-Whitney U and Kruskal - Wallis tests were performed to determine whether there was a statistically significant difference between the number of trade incidents for the various taxa imported and exported. The mean number of exported trades for four periods: 1975–1985, 1986–1996, 1997–2007 and 2008–2018 were also analysed. Poisson regression modelling was used to establish the relationship between zoonotic disease, trade period (years), and the number of trade incidents for the taxa involved. We statistically analysed the data using R (R version 4.0.3, R core team 2020).

## Results

### Species and major countries involved in trade

From 1975–2018, a total of 7,833 trade incidents of wildlife trade were recorded, of which exported and imported incidents from Ghana were 7,084 and 749, respectively. A highly significant difference in the number of exported incidents for the various taxa (Kruskal Wallis test: *p* < 0.01) as well as imported incidents (Kruskal Wallis test: *p* < 0.01) was recorded. Reptiles were the most targeted taxa group, with 81.1 % exports and 70.4 % imports. This was followed by birds with 11 % and 19.2 % exports / imports; then mammals with 7.8 % and 9.3 % export / import; and finally, amphibians with 0.1 % and 1.1 % export / import respectively (Figure 1). The highest number of species classified under CITES Appendix 1 traded belong to the mammal group.

**Figure 1:**
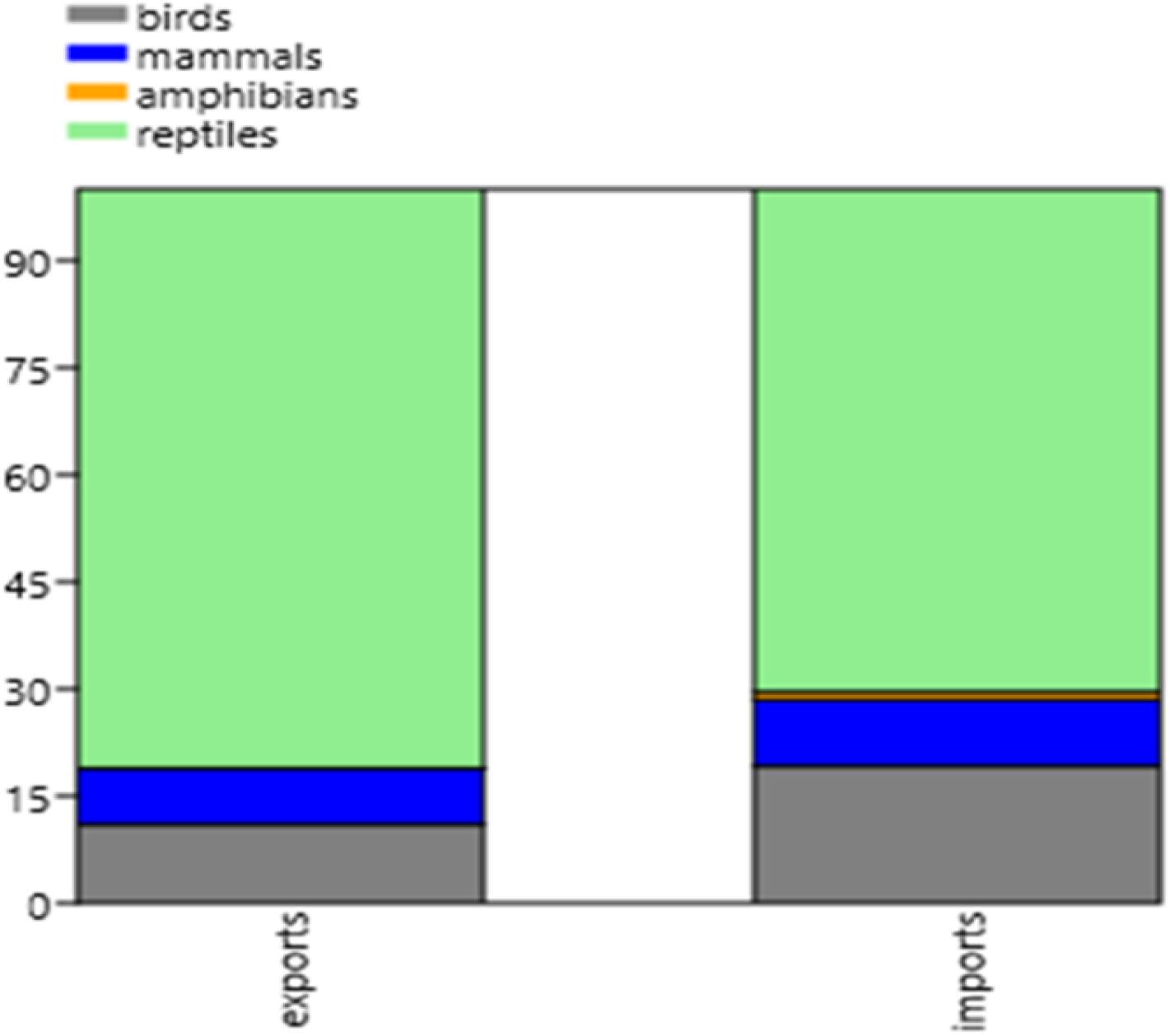
Percentage composition of taxa traded internationally in Ghana

The only amphibian species exported was *Lithobates catesbeianus*. Six species were imported, with *Dendrobates auratus* and *D. leucomelas* (25 % each) as the highest percentage composition (Table 1). A total of 103 bird species accounted for 782 exported incidents from Ghana, and 47 bird species also accounted for 144 incidents imported to Ghana. *Psittacus erithacus* (38.7 % vs 16 %; export vs import) had the highest percentage composition of bird species found in trade, with most traded species constituting the family Psittacidae (Table 1). From 1975–2018, 51 mammal species constituted 552 incidents trade export from Ghana, with *Loxodonta africana*(29.3 %) recording the highest percentage composition (Table 1).

**Table 1:**
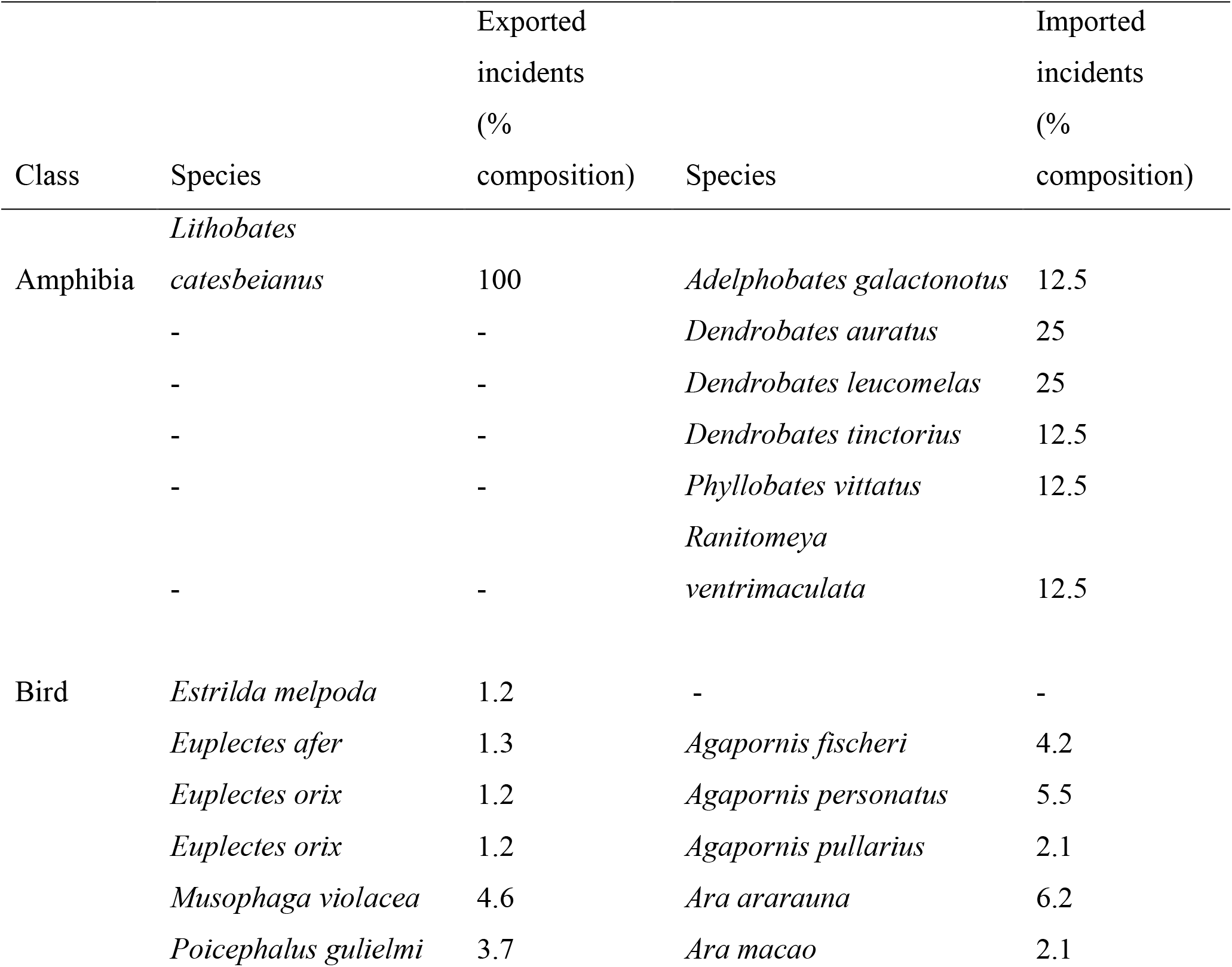
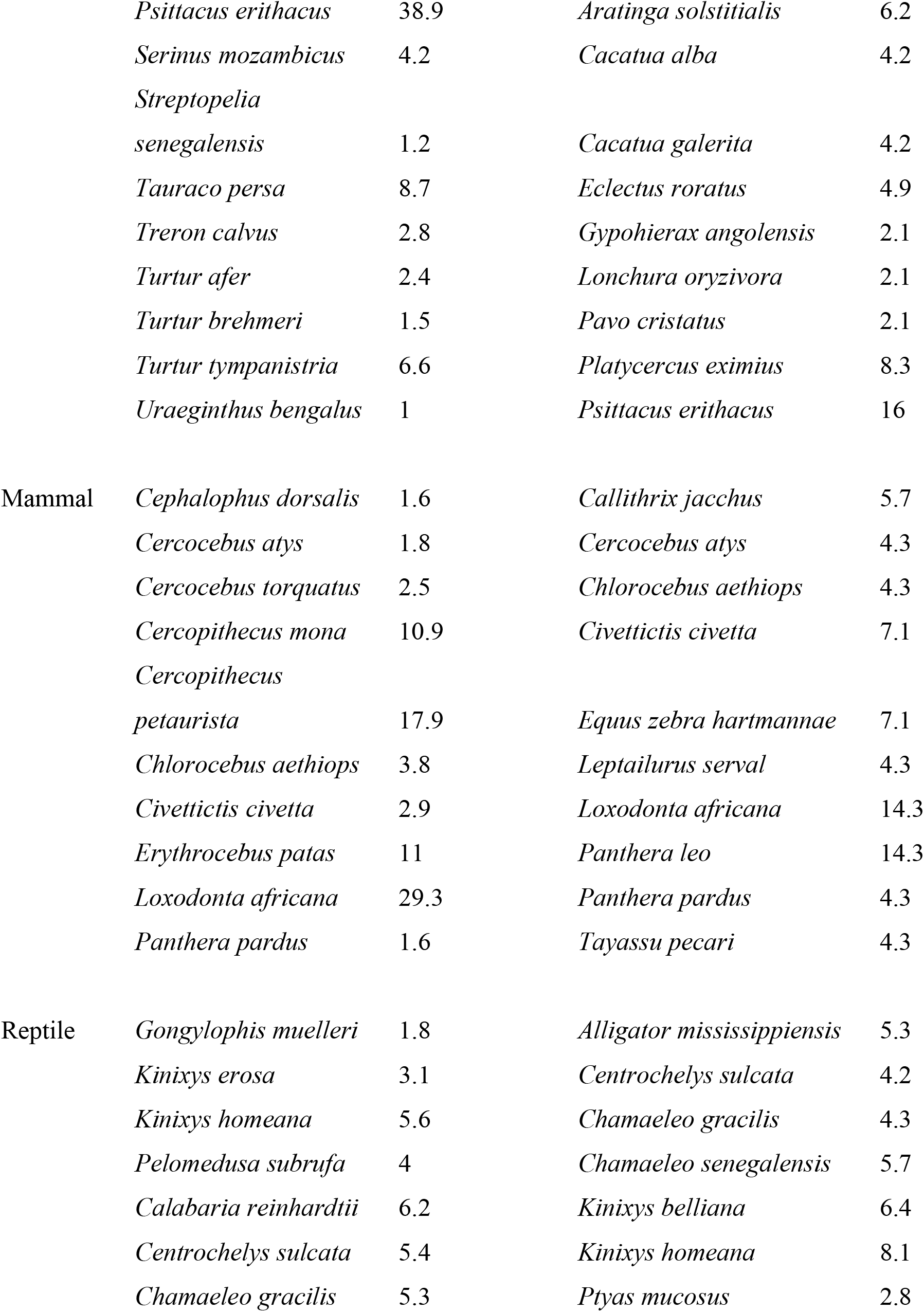
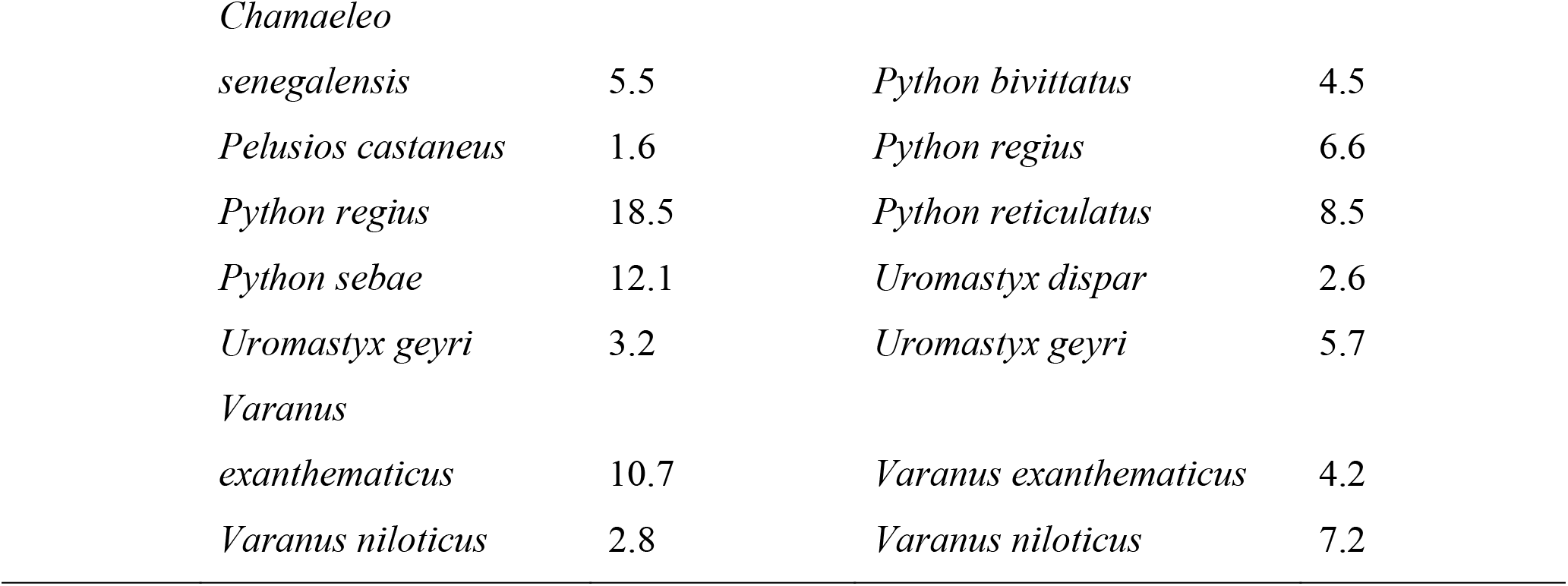
Percentage composition of species for which specimen were traded

| Class | Species | Exported incidents (% composition) | Species | Imported incidents (% composition) |
| --- | --- | --- | --- | --- |
| Amphibia | *Lithobates catesbeianus* | 100 | *Adelphobates galactonotus* | 12.5 |
|  | - | - | *Dendrobates auratus* | 25 |
|  | - | - | *Dendrobates leucomelas* | 25 |
|  | - | - | *Dendrobates tinctorius* | 12.5 |
|  | - | - | *Phyllobates vittatus* | 12.5 |
|  | - | - | *Ranitomeya ventrimaculata* | 12.5 |
|  | - | - | - | - |
| Bird | *Estrilda melpoda* | 1.2 | - | - |
|  | *Euplectes afer* | 1.3 | *Agapornis fischeri* | 4.2 |
|  | *Euplectes orix* | 1.2 | *Agapornis personatus* | 5.5 |
|  | *Euplectes orix* | 1.2 | *Agapornis pullarius* | 2.1 |
|  | *Musophaga violacea* | 4.6 | *Ara ararauna* | 6.2 |
|  | *Poicephalus gulielmi* | 3.7 | *Ara macao* | 2.1 |
|  | *Psittacus erithacus* | 38.9 | *Aratinga solstitialis* | 6.2 |
|  | *Serinus mozambicus* | 4.2 | *Cacatua alba* | 4.2 |
|  | *Streptopelia* |  |  |  |
|  | *senegalensis* | 1.2 | *Cacatua galerita* | 4.2 |
|  | *Tauraco persa* | 8.7 | *Eclectus roratus* | 4.9 |
|  | *Treron calvus* | 2.8 | *Gypohierax angolensis* | 2.1 |
|  | *Turtur afer* | 2.4 | *Lonchura oryzivora* | 2.1 |
|  | *Turtur brehmeri* | 1.5 | *Pavo cristatus* | 2.1 |
|  | *Turtur tympanistria* | 6.6 | *Platycercus eximius* | 8.3 |
|  | *Uraeginthus bengalus* | 1 | *Psittacus erithacus* | 16 |
| Mammal | *Cephalophus dorsalis* | 1.6 | *Callithrix jacchus* | 5.7 |
|  | *Cercocebus atys* | 1.8 | *Cercocebus atys* | 4.3 |
|  | *Cercocebus torquatus* | 2.5 | *Chlorocebus aethiops* | 4.3 |
|  | *Cercopithecus mona* | 10.9 | *Civettictis civetta* | 7.1 |
|  | *Cercopithecus* |  |  |  |
|  | *petaurista* | 17.9 | *Equus zebra hartmannae* | 7.1 |
|  | *Chlorocebus aethiops* | 3.8 | *Leptailurus serval* | 4.3 |
|  | *Civettictis civetta* | 2.9 | *Loxodonta africana* | 14.3 |
|  | *Erythrocebus patas* | 11 | *Panthera leo* | 14.3 |
|  | *Loxodonta africana* | 29.3 | *Panthera pardus* | 4.3 |
|  | *Panthera pardus* | 1.6 | *Tayassu pecari* | 4.3 |
| Reptile | *Gongylophis muelleri* | 1.8 | *Alligator mississippiensis* | 5.3 |
|  | *Kinixys erosa* | 3.1 | *Centrochelys sulcata* | 4.2 |
|  | *Kinixys homeana* | 5.6 | *Chamaeleo gracilis* | 4.3 |
|  | *Pelomedusa subrufa* | 4 | *Chamaeleo senegalensis* | 5.7 |
|  | *Calabaria reinhardtii* | 6.2 | *Kinixys belliana* | 6.4 |
|  | *Centrochelys sulcata* | 5.4 | *Kinixys homeana* | 8.1 |
|  | *Chamaeleo gracilis* | 5.3 | *Ptyas mucosus* | 2.8 |

|  |  |  |  |
| --- | --- | --- | --- |
| *Chamaeleo* |  |  |  |
| *senegalensis* | 5.5 | *Python bivittatus* | 4.5 |
| *Pelusios castaneus* | 1.6 | *Python regius* | 6.6 |
| *Python regius* | 18.5 | *Python reticulatus* | 8.5 |
| *Python sebae* | 12.1 | *Uromastix dispar* | 2.6 |
| *Uromastix geyri* | 3.2 | *Uromastix geyri* | 5.7 |
| *Varanus* |  |  |  |
| *exanthematicus* | 10.7 | *Varanus exanthematicus* | 4.2 |
| *Varanus niloticus* | 2.8 | *Varanus niloticus* | 7.2 |

*Panthera leo* and *Loxodonta africana* (14.3 % each) constituted the highest percentage composition among 29 mammal species imported to Ghana. The majority of mammal species traded belonged to the family Cercopithecidae for export and Felidae (imports). Among the 71 reptile species involved in exported incidents from Ghana, *Python regius* (18.5 %) had the highest percentage composition, while *Python reticulatus* (8.3 %) recorded the highest import among the 49 reptile species. Reptile species of the family Pythonidae were the most traded (Table 1). Mann – Whitney U test indicated significant differences in the number of taxa trade between imported and exported incidents for birds (U = 2503.5,*p* < 0.05), mammals (U = 610.5,*p* < 0.05) and reptiles (U = 1545.5,*p* < 0.05).

Major countries involved in wildlife trade with Ghana and highly linked taxa include but are not limited to Germany (amphibians); Japan, Italy and South Africa (birds); United States of America (USA), United Kingdom (UK) and South Africa (mammals) and the USA, Japan and Benin (reptiles, Table 2). Most importer countries are from Europe and the USA, while countries that export wildlife to Ghana are from Africa.

**Table 2:** Countries with the highest number of trade incidents

| Taxa | Importer country | No. of trade incidents | Exporter country | No. of trade incidents |
| --- | --- | --- | --- | --- |
| Amphibian | Czech Republic | 1 | Germany | 8 |
|  | UK | 1 |  |  |
| Bird | Japan | 91 | South Africa | 77 |
|  | Italy | 87 | Netherlands | 26 |
|  | USA | 78 | Indonesia | 13 |
|  | UK | 59 | Togo | 6 |
| Mammal | USA | 157 | South Africa | 14 |
|  | UK | 63 | Zimbabwe | 10 |
|  | Japan | 40 | Namibia | 5 |
|  | Germany | 32 | Botswana | 4 |
| Reptile | USA | 1369 | Benin | 136 |
|  | Japan | 492 | Togo | 87 |
|  | UK | 489 | Mali | 64 |
|  | Germany | 331 | Indonesia | 40 |

### Trends in the period of trade and zoonotic diseases

The international wildlife trade of Ghana varied from year to year (Figure 2 and 3). The highest number of trade for taxa included: birds (n=52, export) and (n=26, import) recorded in the year 2004 and 2012; mammals (n=39, export) and (n=8, import) recorded in the year 1986 and 2014; reptiles (n=268, export) and (n=50, import) recorded in the year 2004 and 2012 respectively. It is important to note that there was a decrease in mammal trade in the years in which birds and reptiles had the highest number of trades. The mean number of exported trades for four periods: 1975–1985, 1986–1996, 1997–2007 and 2008–2018 were analysed. The decade mean number of trade for 1997–2007 was the highest (62 ± 13.69) followed by 2008–2018 (54.4 ± 13.45), then 1986–1996 (46.9 ± 8.81) and the least was in 1975–1985 (14.7 ± 2.32).

**Figure 2:**
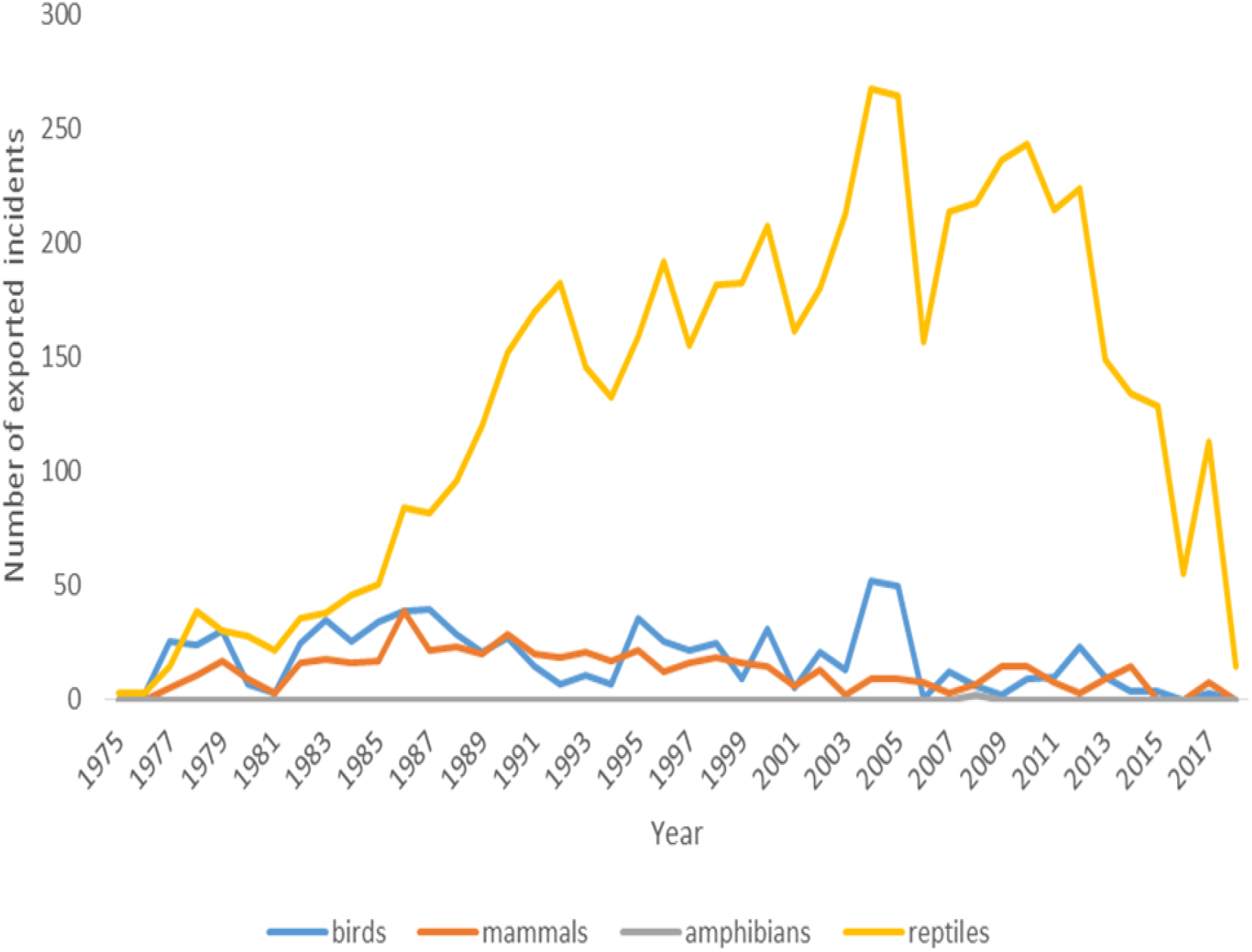
Trend in the number of exported incidents from 1975–2018

**Figure 3:**
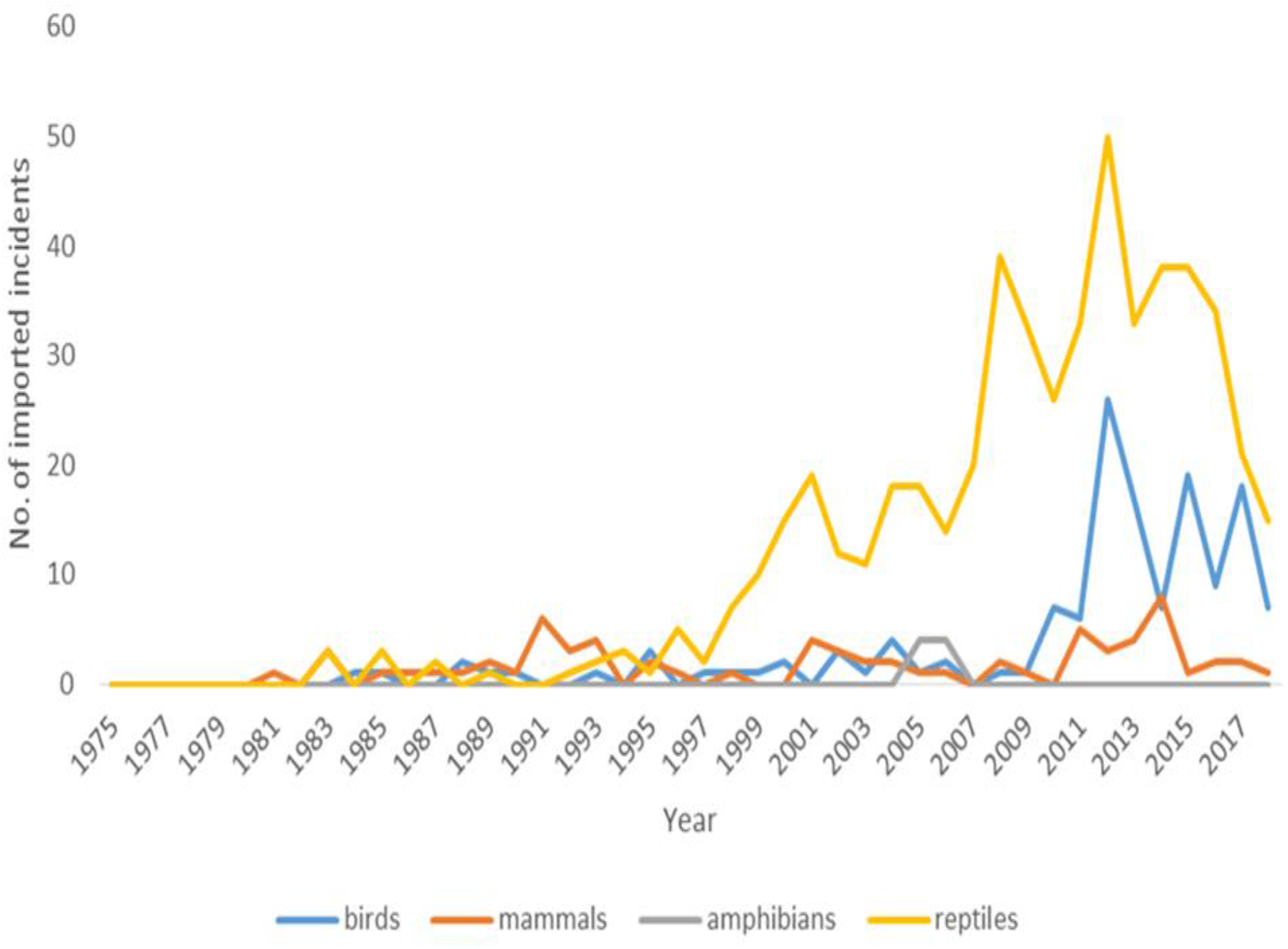
Trend in the number of imported incidents from 1975–2018

A significant negative relationship between years of trade and disease outbreaks and mammal trade was found. Trade in reptiles showed a significant positive relationship with years of trade and disease outbreaks (Table 3). Suggesting that mammal trade decreased over the years and with an outbreak of disease, while reptile trade increased. Table 3 showed a significant negative relationship between disease outbreak and bird import (*Coef*. = −0.56, *SE* = 0.193, *Z*-value = −2.9, *p* = 0.004, Table 3).

**Table 3:** The generalised linear model results for relationships between taxa specimen traded (export and import) and explanatory variables.

| Taxa | Variables | Export |  |  |  | Import |  |  |  |
| --- | --- | --- | --- | --- | --- | --- | --- | --- | --- |
|  |  | Coef. | SE | z-value | p-value | Coef. | SE | z-value | p-value |
| Amphibians | Years | 0.08 | 0.07 | 1.13 | 0.258 | 0.06 | 0.033 | 1.81 | 0.07 |
|  | Disease outbreak | -13.7 | 2553.13 | -0.01 | 0.996 | -14.68 | 2055.7 | -0.01 | 0.994 |
| Birds | Years | -0.01 | 0.003 | -6.86 | < 0.001 | 0.13 | 0.012 | 11.22 | < 0.001 |
|  | Disease outbreak | 0.49 | 0.072 | 6.94 | < 0.001 | -0.56 | 0.193 | -2.9 | 0.004 |
| Mammals | Years | -0.12 | 0.004 | -3.24 | 0.001 | 0.04 | 0.011 | 3.52 | < 0.001 |
|  | Disease outbreak | -0.45 | 0.091 | -4.99 | < 0.001 | -0.19 | 0.26 | -0.75 | 0.45 |
| Reptiles | Years | 0.03 | 0.001 | 26.97 | < 0.001 | 0.10 | 0.005 | 19.82 | < 0.001 |
|  | Disease outbreak | 0.30 | 0.044 | 6.94 | < 0.001 | 0.56 | 0.131 | 4.3 | < 0.001 |

## Discussion

The alarming rate at which wildlife specimens are traded with associated problems of poaching and infectious disease emergence has become a global concern, especially for species threatened with extinction whose rareness increases exploitation. We found that mammals were the most negatively significantly affected with the outbreak of zoonotic diseases while trade in reptiles increased during the same period. Imports of bird specimen had a negatively significant relationship with zoonotic disease outbreak. Most exporter countries are from Africa and importers from the USA and Europe. Wildlife trade in Ghana withnessed significant exports than imports targeting the most threatened species including *L. Africana*, psitaccus and pthypons. The highest number of trade occurred in the decade 1997–2007 and 2008–2018.

The exploitation of wildlife for trade is no different in Ghana. Ghana had high incidents of export of wildlife specimens (90.44 %) but saw low import incidents (9.56 %). This is consistent with trade in Australia and Mexico, both with high export and low import (Wyatt 2016; Arroyo-Quiroz and Wyatt 2019). Contrary to our study, New Zealand and the UK have high import and low export incidents (Wyatt 2016). This probably could be that countries with high export are rich in biodiversity. This explains why Ghana, Mexico, and Australia have high export. Mexico and Australia are part of the mega-biodiversity countries globally (Brooks et al. 2006). While importer countries have the financial strength to buy wildlife specimen considered as sought-after symbol of stature and wealth. A study by Scheffers et al. (2019) found that some countries have high trade links to specific taxa, confirming the results of our study. The USA, Japan and European countries highly imported reptiles from Ghana and this is consistent with other studies (Arroyo-Quiroz and Wyatt 2019)

In this study, reptiles constituted the majority in demand for international trade, followed by birds. This is consistent with other wildlife trade studies (Robinson et al. 2015; Alacs and Georges 2008). However, a global study of wildlife trade found reptiles the least traded and birds most traded (Scheffers et al. 2019). This suggests a mixed result compared to our study. A high reptile and bird trade incident found in our study could be attributed to ease in handling, high diversity, and availability of the market for pets (Alacs and Georges 2008; Fukushima et al. 2020). Traders favour reptiles because they derive substantial prices on the market when sold as pets. Interestingly, we observed the least number of incidents for amphibians. This might contradict the reasoning of ease of handling for high trade.

Most wildlife species were targeted for body parts such as skin and trophies for artefacts and used as pets, food, and medical experiments (Nijman 2010; Rosen and Smith 2010; de Magalhães and São-Pedro 2012) as the underlying reason for high trade in python species, *Psittacus erithacus*, and *L. africana.* Most python species are threatened for the exploitation of their skin because of their large size (Natusch and Lyons 2014; Luiselli and Akani 2002) and the pet trade (Bartlett et al., 2001). This also could explain why amphibians were least traded due to low demand to satisfy the majority of the reasons aforementioned. Again, many amphibians are common; traders, for example, will not spend billions of dollars on easily accessible species. Low trade incidents in amphibians could also be that the majority of amphibians traded were not detected or undeclared. The survival of *L. africana* is threatened by an increased rate of poaching purposely for its ivory.

Items made of ivory are in high demand for high prices as it indicates the symbol of stature and wealth of the bearer in some societies, therefore has a high purchasing power of the consumer (CITES 2012). It is suggested that the ivory has medicinal properties to cure diseases, especially cancer, hence the high supply of the specimens (Anderson and Jooste 2014). The only amphibian species exported from Ghana was *L. catesbeianus*. This frog is native to eastern North America. There are no indications of its presence in sub-Saharan African. It is very clear from this that Ghana might have been used as a transit for this species to its destination. *Psittacus erithacus* accounted for the largest proportion of birds traded in Ghana. This is consistent with a study in China by Li and Jiang (2014). In the circles of trade, it is among the most highly prized bird for its human speech mimicking capabilities and its intelligence (Annorbah et al. 2016).

The impact of disease outbreaks varied among taxa. Mammals were the most negatively affected. Both export and import of mammal specimens decreased significantly with the outbreak of diseases. Mammals are found to be significantly probable to host zoonotic pathogens, with primates and bats sharing the greatest proportion as hosts (Han et al. 2016; Johnson et al. 2020). Many of the high risk/severe infectious diseases are linked to mammals. Few examples include Ebola virus (primates), monkeypox (African rodents), HIV (primates), SARS (bats, Bell et al. 2004; Leroy et al. 2004) and recent speculation of Coronavirus (COVID-19) in bats and pangolin (Rajgor et al. 2020; Zhang et al. 2020).

The reduction in mammal trade over the years with the outbreak of diseases could be attributed to two reasons; first, most of these diseases become global or regional pandemics that require regulatory bodies to raise awareness on trade in such animal groups. For example, the World Health Organization (WHO) places a “Disease X” and classifies it as a disease requiring high alert, and therefore advocates policies against any form of animal-human contact (Borsky et al. 2020).

In addition, a ban on all forms of trade relating to a specific taxonomic group is carried out in some instances (e.g., China, Zhou et al. 2020). Second, such diseases’ severity and death potentials cause fear and facilitate a shift by traders from animals potentially hosting such diseases.

Fear of industry players being infected with diseases and subsequent deaths makes them consciously shift their trade activities to other wild animals with records of no or little disease infection impacts over some time. For example, mammal trade showed a sharp decline from 2015–2018, the same period for the Ebola disease outbreak, which caused devastating effects to human health and economies. Similar events occurred from 2003–2007, following the outbreak of SARS disease. The same can be mentioned for bird trade decline during 2006–2011, following the avian influenza disease outbreak. During these periods where mammal and bird trade decreased, reptile trade, on the contrary, increased. It could be deduced that traders shifted their activities among mammals, birds, and reptiles with the outbreaks of those diseases. In the years 2004 and 2012, birds and reptiles recorded the highest number of trades. However, mammals’ trade decreased significantly, and these periods also witnessed the outbreaks of highly transmittable diseases, such as Middle East Respiratory Syndrome (MERS). An outbreak of zoonotic diseases is likely to cause shifts in taxa trade globally.

## Conclusion and Recommendations

Wildlife is under intense pressure from trade as thousands of incidents of CITES-listed species of wildlife are found in trade. Reptiles were highly affected in trade both for export and import, followed by birds, mammals, and amphibians, respectively. The highest number of trade flows occurred in 1997-2007. Continuous trade in reptiles and birds, especially the endangered pythons and Psittacus species, could lead to their extinction in the wild. The outbreak of zoonotic diseases influences demands for different taxa at different periods, hence the dynamics of international wildlife trade in Ghana largely through exports. The outbreak of diseases caused a decline in mammal trade while reptile trade increased over the years. Early detection of zoonotic diseases and the adoption of an expanded education module on avoiding species capable of harbouring pathogens will most likely help reduce trade in wildlife. Furthermore, considerable cooperation among countries in implementing wildlife trade regulations will also help to reduce wildlife trade impacts.

## Acknowledgement

The data source for this study was derived from the CITES Trade Database: “CITES trade statistics derived from the CITES Trade Database, UNEP World Conservation Monitoring Centre, Cambridge, UK.”

## Data availability statement

All numbers used in the analyses were generated from the Convention on Intentional Trade in Endangered Species of Wild Fauna and Flora (CITES) database. Accessed on 12 October 2019 from http://www.unep-wcmc.org/citestrade

## Declaration

The authors declare that the paper sent is original, and no part of it has been published before or is being considered for publication in any other journal.

